# Immunosequencing reveals diagnostic signatures of chronic viral infection in T cell memory

**DOI:** 10.1101/026567

**Authors:** Ryan O. Emerson, William S. DeWitt, Marissa Vignali, Jenna Gravley, Cindy Desmarais, Christopher S. Carlson, John A. Hansen, Mark Rieder, Harlan S. Robins

## Abstract

B and T cells expand clonally in response to pathogenic infection, and their descendants, which share the same receptor sequence, can persist for years, forming the basis of immunological memory. While most T cell receptor (TCR) sequences are seen very rarely, ‘public’ TCRs are present in many individuals.

Using a combination of high throughput immunosequencing, statistical association of particular TCRs with disease status, and machine learning, we identified of a set of public TCRs that discriminates cytomegalovirus (CMV) infection status with high accuracy. This pathogen-specific diagnostic tool uses a very general assay that relies only on a training cohort coupled with immunosequencing and sophisticated data analysis. Since all memory T cell responses are encoded in the common format of somatic TCR rearrangements, a key advantage of reading T cell memory to predict disease status is that this approach should apply to a wide variety of diseases. The underlying dataset is the largest collection of TCRs ever published, including ∼300 gigabases of sequencing data and ∼85 million unique TCRs across 640 HLA-typed individuals, which constitutes by far the largest such collection ever published. We expect these data to be a valuable public resource for researchers studying the TCR repertoire.

## INTRODUCTION

The ability of the cellular adaptive immune system to adequately address an incipient infection relies on the presence of B and T cells that have generated appropriate antigen-specific receptors. Upon antigen recognition, activated T cells proliferate by clonal expansion and become part of the memory T cell compartment, where they reside for many years as a clonal population of cells with identical-by-descent rearranged T cell receptor (TCR) genes ^1^^-^^3^.

The majority of the TCR diversity resides in the β chain of the αβ heterodimeric TCR. Each mature TCR_β_ gene is randomly rearranged from the germ-line locus by combining noncontiguous TCR_β_ variable (V), diversity (D), and joining (J) region gene segments, which collectively encode the CDR3 region (the primary region of the TCRβ locus for determining antigen specificity). Deletion and template-independent insertion of nucleotides during rearrangement at the V_β_-D_β_ and D_β_-J_β_ junctions further add to the potential diversity of receptors that can be encoded ^4^^,^^5^.

The interaction of TCRs with their cognate antigen is mediated by the cell-surface presentation of foreign peptides by pathogen-infected cells in the context of major histocompatibility complex (MHC) class I proteins. Since MHC class I proteins are encoded by the human leukocyte antigen (HLA) loci A, B, and C, which are highly polymorphic, the antigen specificity of a TCR is modulated across individuals by HLA context.

Healthy adults express approximately 10 million unique TCRβ chains on their 10^12^ circulating T cells^3^. Despite the fact that these are drawn from a much larger pool of possible rearrangements, observing the same TCRβ chain independently in two individuals is thousands of times more common than would be expected if all rearrangements were equally likely ^6^. Therefore, it is expected that many specific TCRβ sequences (especially those with few or no junctional insertions) are present in the naïve T cell repertoires of most humans at any given time and will reliably proliferate upon exposure to their target antigen in the proper MHC context ^7^. This over-representation of specific TCRβ sequence rearrangements in the naïve T cell repertoire forms the basis of public T cell responses, in which a particular antigen is targeted by the same T cell receptor sequence in multiple individuals ^8^^,^^9^. Public T cell responses are observed when the space of potential high-avidity TCRβ chains that could bind to a particular antigen-MHC complex includes one or more TCRβ chains that also have a high likelihood of existing in the naïve repertoire at any given time. Sequences associated with a public T cell response to a particular antigen will only be intermittently present in the naïve compartment of subjects that have not been exposed to that antigen. However, such TCRβ sequences should consistently appear in the T cell repertoire of subjects who have been exposed to the antigen, having undergone clonal expansion after antigen encounter, and providing a basis for comparing immunological memory across different individuals. Despite historical limitations on sequencing depth and the limited size of investigational cohorts, previous work has identified many individual examples of public T cell responses to infectious diseases (including CMV, EBV, *C. tetani*, parvovirus, HSV, HIV and influenza) as well as in malignancies and autoimmunity ^8^^,^^10^. Typically these public T cell responses have been studied in the context of single antigens in a single HLA context, usually using purified antigen-MHC complexes and fluorescent tagging to isolate antigen-specific T cells.

In this study, we aimed to develop a diagnostic strategy that is highly specific for a particular disease status, while using data from a very general assay that relies only on a training cohort coupled with immunosequencing and sophisticated data analysis. As a first step to identify significant associations between sets of TCRβ sequences and a certain disease status, we measured millions of distinct traits (i.e. the presence and abundance of T cell receptor sequences) in a large investigational cohort and statistically assessed the concordance of each such trait with a phenotype of interest. For our proof-of-principle experiment we selected cytomegalovirus (CMV) infection as the phenotype of interest. CMV results in a chronic viral infection, is present in 30-90% of adults depending on the population studied ^11^, and has been extensively studied as a model system for public T cell responses.

## RESULTS

### Identification of CMV-associated TCRβ sequences

Our strategy, as illustrated in Figure 1, began with the high-throughput characterization of rearranged TCR genes in 640 healthy subjects with known CMV status. Subject demographics are presented in Table 1, and the resulting immunosequencing data are publicly available at http://adaptivebiotech.com/pub/emerson-2015. Approximately 185,000 different TCRβ sequences were observed per subject, each presumably specific for an antigen:MHC complex of unknown identity. We then searched for TCRβ sequences (i.e., TCRβ DNA sequences implying identical putative full-length TCRβ proteins) present in multiple subjects, and identified a set of TCRβ sequences that were significantly associated with positive CMV status. Briefly, we calculated a *P* value for the association of each TCRβ sequence with CMV status using a Fisher exact test, controlling the false discovery rate (FDR) by permutation of the CMV status (see Materials and Methods), and we identified a list of CMV-associated TCRβ sequences (for a certain FDR and *P* value, see below). Next, we calculated a CMV memory burden for each subject as the proportion of all that subject’s TCRβ sequences that are represented in the catalog of CMV-associated TCRβ sequences. Finally, we attempted to use this CMV memory burden to distinguish between CMV+ and CMV- subjects.

**Figure 1:**
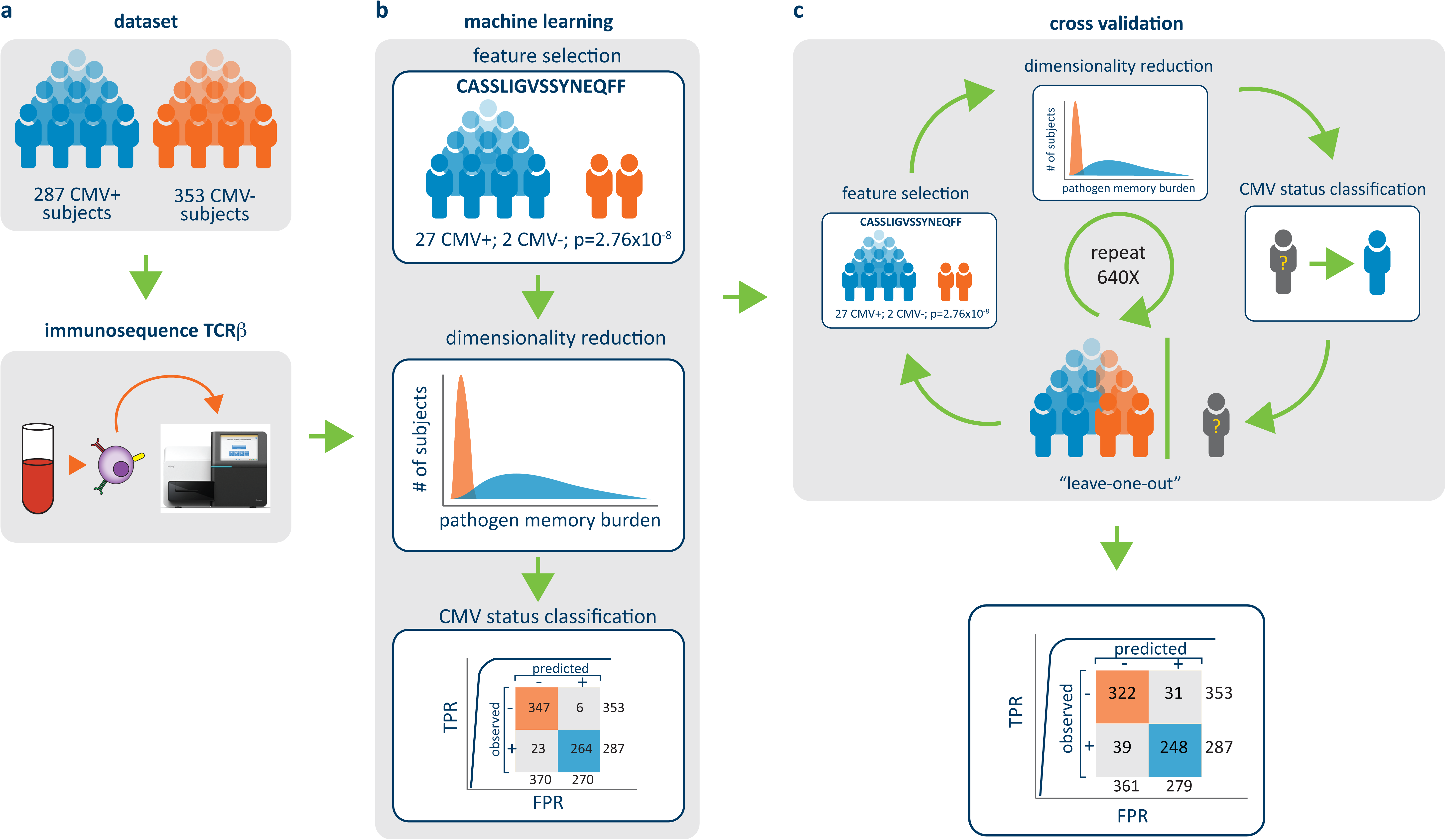
Experimental and analytical flow. **(a)** We analyzed peripheral blood samples from 640 healthy subjects (287 CMV- and 353 CMV+) by high-throughput TCRβ profiling. **(b)** We identified TCRβ sequences that were present in significantly more CMV+ subjects than CMV- subjects, controlling FDR by permutation of CMV status. These data were used to build a classification model. **(c)** The model was tested using exhaustive leave-one-out cross-validation, in which one sample was held out and the process repeated from the beginning. The resulting classification model was used to predict the CMV status of the holdout subject.

**Table 1:** Cohort demographics. Age, sex, race/ethnicity and CMV status for the 640 subjects in our study cohort.

| **Sex** | **CMV+** | **CMV-** | **All** |
| --- | --- | --- | --- |
| Female | 150 | 147 | **297** |
| Male | 137 | 206 | **343** |
| **All** | **287** | **353** | **640** |

| **Race/ethnicity** | **CMV+** | **CMV-** | **All** |
| --- | --- | --- | --- |
| White | 164 | 212 | **376** |
| Black or African American | 8 | 0 | **8** |
| Asian | 15 | 2 | **17** |
| American Indian or Alaska Native | 7 | 2 | **9** |
| Native Hawaiian or other Pacific Islander | 3 | 0 | **3** |
| Hispanic or Latino | 20 | 6 | **26** |
| Unknown | 70 | 131 | **201** |
| **All** | **287** | **353** | **640** |

\* NIH categories used, all Hispanic/Latino are classified as race = Unknown

| **Age (in years)** | **CMV+** | **CMV-** | **All** |
| --- | --- | --- | --- |
| Mean | 42 | 37 | 40 |
| Median | 44 | 38 | 41 |
| Range | 5-74 | 1-70 | 1-74 |
| # of patients of unknown age | 31 | 56 | 87 |

Figure 2 presents the results of our machine learning approach: determining the CMV-associated TCRs and the CMV memory burden as described above, we performed a logistic regression to separate CMV+ from CMV- subjects by CMV memory burden. To test the robustness of our results, we varied the p-value threshold for inclusion in the catalog of CMV-associated TCRβ sequences. Fig. 2A shows the performance (as measured by the area under the ROC curve, or AUROC) for both the full dataset and an exhaustive leave-one-out cross-validation dataset, and Fig2B shows the estimated FDR for a range of p-value thresholds. The best performance is seen at a *P* value of 10^-4^, which corresponds to an estimated FDR of ∼20%, resulting in the identification of a set of 142 TCRβ sequences that were significantly associated with positive CMV status (listed in Supplemental Table 1). Using these conditions results in good separation between the CMV+ and CMV- subjects in our cohort as measured by CMV memory burden (Figure 2C). Finally, Figure 2D shows the ROC curves for both the full and the cross-validation datasets. The AUROC for the full dataset is 0.98, indicating that our approach resulted in an excellent classifier for CMV status. In addition, at the point of highest discriminating power, we observe an accuracy of 0.89 and a diagnostic odds ratio of 66 in the cross-validation dataset. Taken together, these data suggest that that presence of public T cell responses to CMV is highly correlated with CMV positive status.

**Figure 2:**
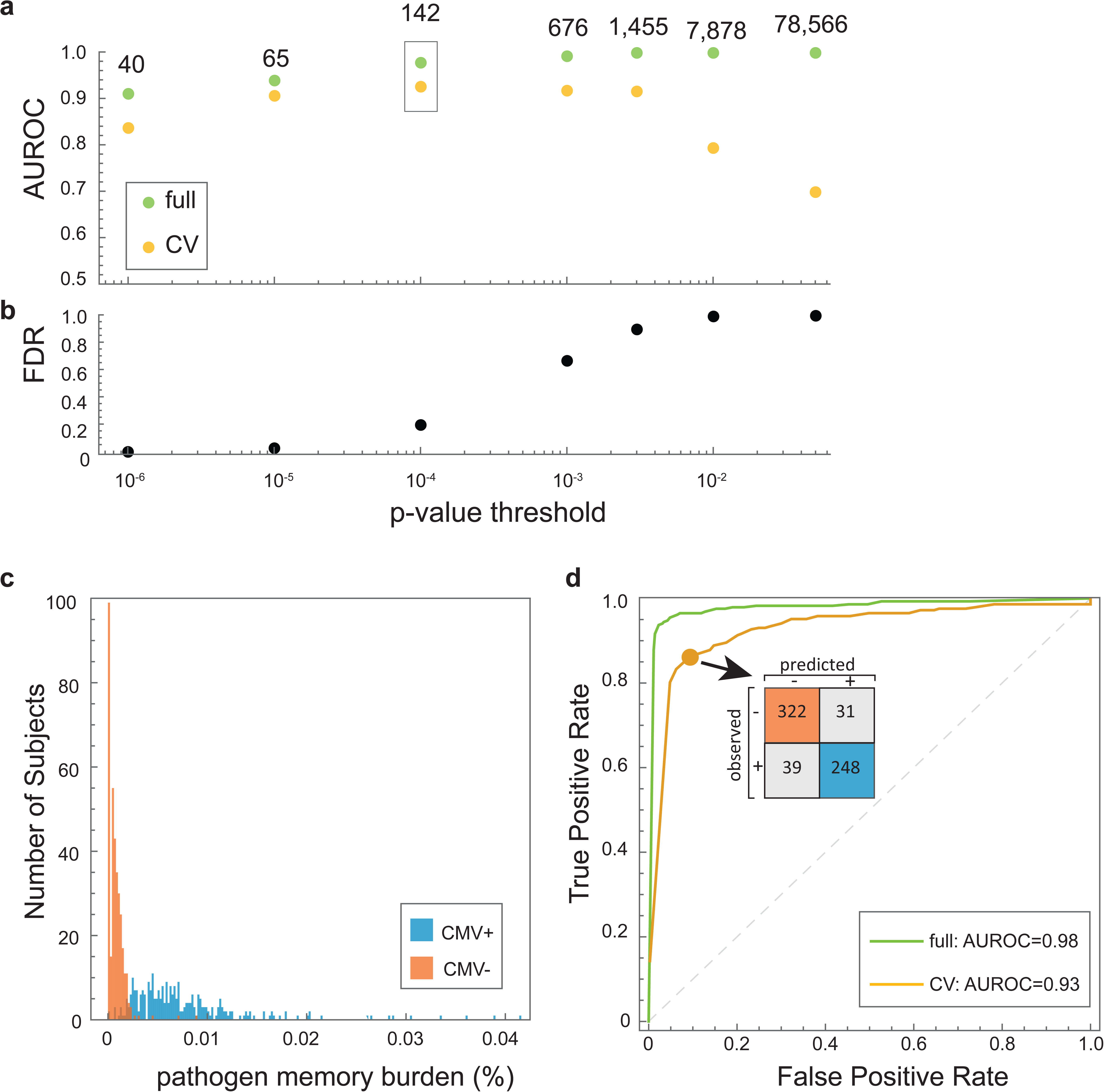
Machine learning results. **(a)** Classification performance on the full (•) and cross-validation (CV) (•) datasets for each p-value threshold, measured as the area under the ROC curve (AUROC). The numbers correspond to the CMV-associated TCRβ identified at each p-value threshold; we selected the dataset corresponding to p ≤ 1×10^-4^ (boxed) for downstream analyses. **(b)** False discovery rate (FDR) estimated for each p-value threshold used in the identification of significantly CMV-associated TCRβ sequences, using permutations of CMV status. **(c)** Distribution of CMV memory burden (i.e., the proportion of each subject’s TCRβ repertoire that matches our list of 142 CMV-associated TCRβ sequences) among CMV+ and CMV- subjects. **(d)** ROC curves calculated for the full and cross-validation datasets, and confusion matrix calculated on the cross-validation dataset. The highest accuracy (0.89, diagnostic odds ratio 66) is achieved when classifying 86% of true positives correctly with a false positive rate of 8%.

### HLA association analysis

Given that T cells recognize their cognate antigens in the context of MHC molecules expressed by antigen presenting cells, next we wanted to test whether we could identify the HLA-restriction of our CMV-associated TCRβ sequences. We performed a Fisher’s exact test on each CMV-associated TCRβ sequence to determine if its presence was significantly associated with any of the HLA alleles observed in the cohort. We could confidently assign the association of 57 out of 142 CMV-associated TCRβ sequences with at least one HLA allele, with a p-value cutoff of 1×10^-3^. Full results are presented in Figure 3 and Supplemental Table 1.

**Figure 3:**
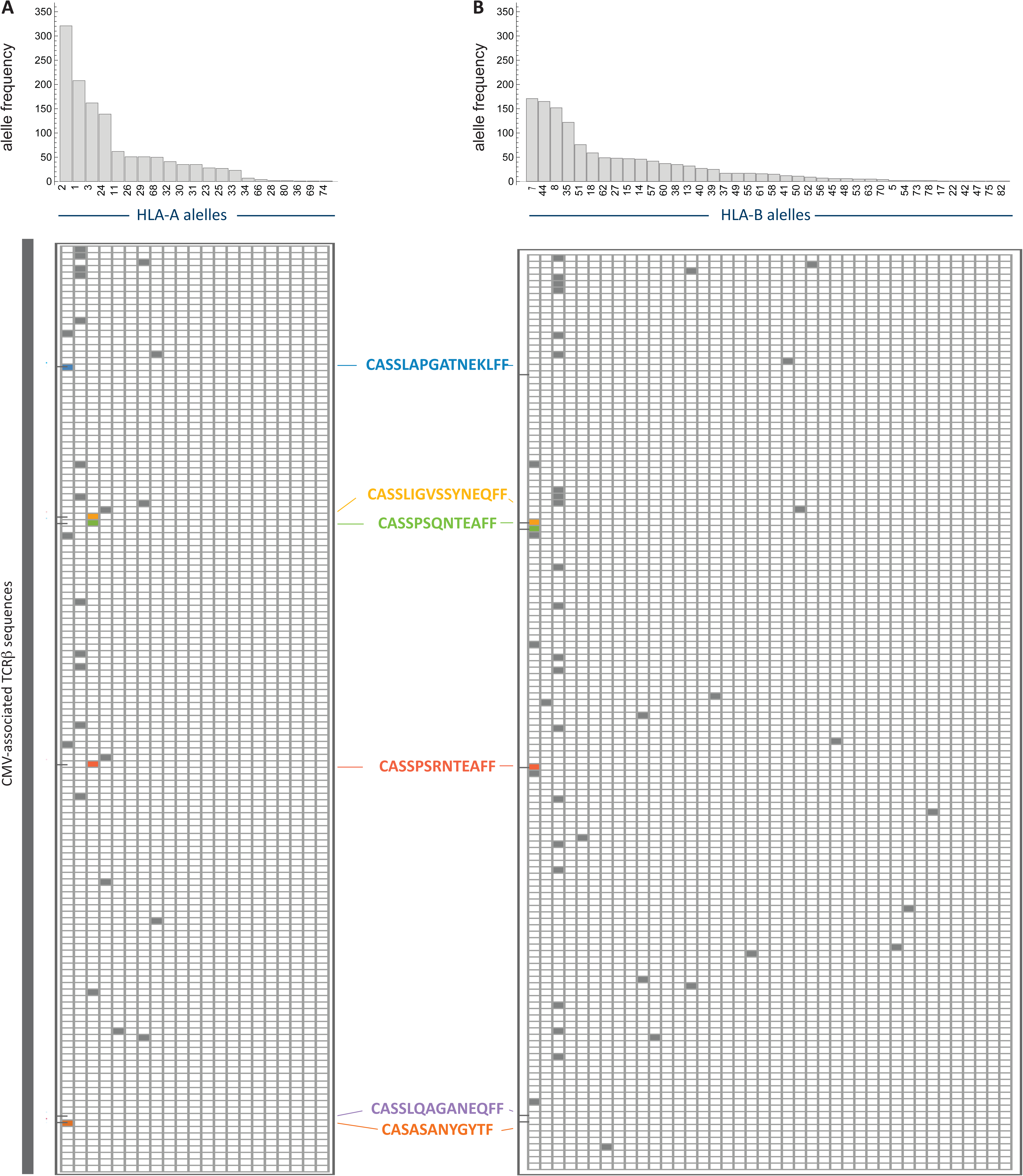
HLA-restriction of CMV-associated TCRβ sequences. Distribution of HLA-A **(a)** and HLA-B **(b)** alleles in this cohort. **(c)** Each of the 142 CMV-associated TCRβ sequences (p ≤ 1×10^-4^) was tested for significant association with each HLA allele, with a p-value threshold of 1×10^-3^. Of these, 57 are significantly associated with an HLA-A and/or an HLA-B allele, and none are significantly associated with more than a single allele from each locus. The colored sequences correspond to those that had been previously identified as CMV-associated (see Table 2); in 4 cases we recapitulate the correct HLA association, while we did not see a statistically significant HLA association for the remaining sequence.

### Comparison to previously identified CMV-associated TCRβ sequences

Finally, we performed a literature search to identify previously reported CMV-reactive TCRβ sequences. We found 595 unique TCRβ sequences that had been reported by at least one previously published study ^7^^,^^10^^,^^12^^-^^27^. Many of these are observed in our dataset, but most are seen in roughly equal number of CMV+ and CMV- subjects (Figure 4). This observation could be explained by receptor sequences with exceptionally high frequency in the naïve repertoire, or could reflect cross-reactive receptors that bind to CMV antigens but also other common antigens. Of these unique TCRβ sequences, 565 were detected in one individual in a single study, whereas 30 had previously been classified as public (i.e., seen in multiple individuals in one study, on in multiple studies). Moreover, the public TCRβ sequences reported in the literature are considerably more common in our cohort than those previously identified in a single individual.

**Figure 4:**
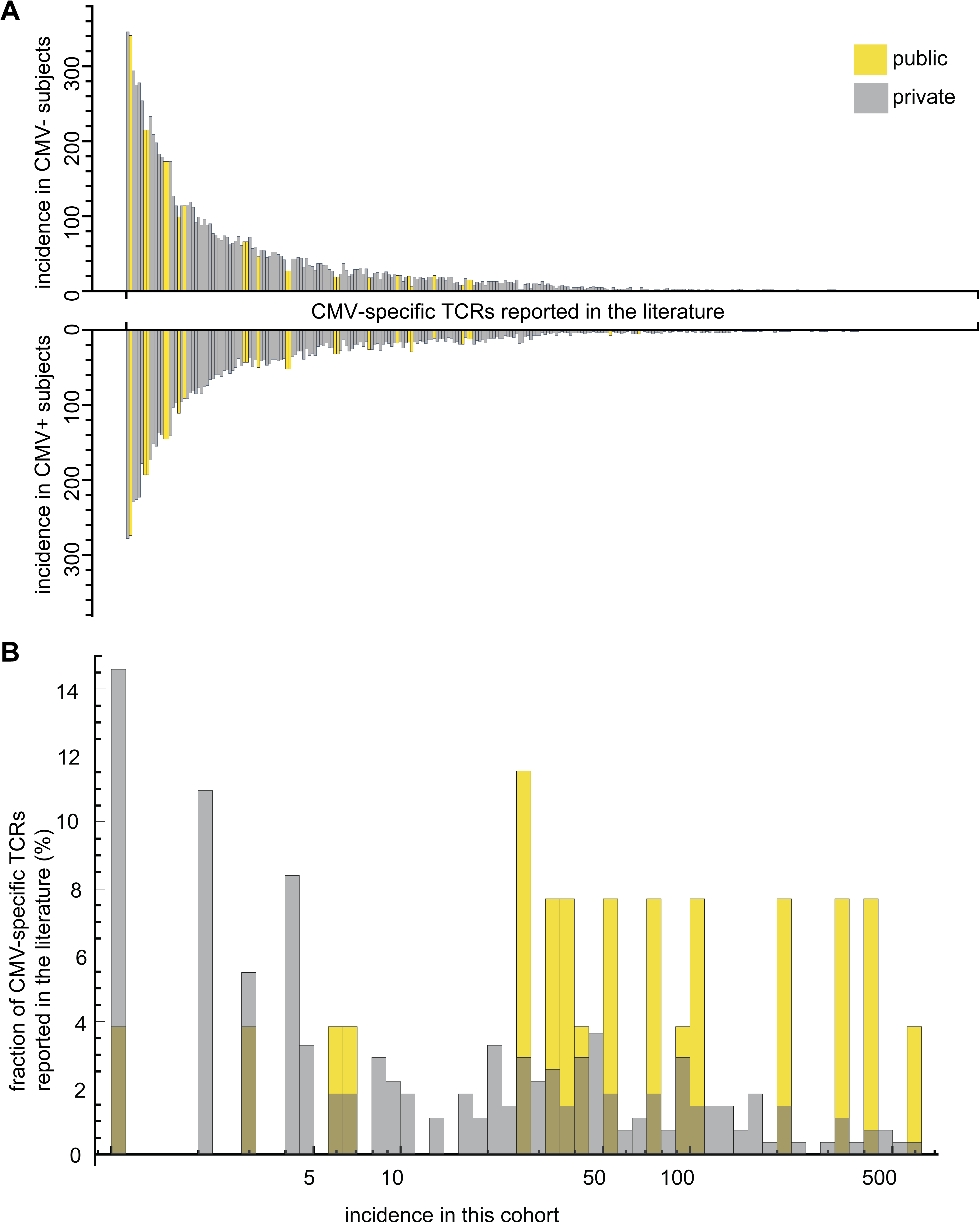
Incidence of previously reported CMV-reactive TCRβ sequences in this dataset. **(a)** The incidence of each previously published CMV-associated TCRβ sequences in our cohort is plotted along the horizontal axis by decreasing total incidence (CMV+ subjects above the horizontal and CMV- subjects below the horizontal line). **(b)** The incidence of these TCRβ sequences in our cohort of 640 subjects for each group of sequences is shown.

Six of these 30 public TCRβ sequences were contained in our set of 142 CMV- associated TCRβ sequences (Table 2). The concordance between the V and J genes we report and previously published studies is very good, although previously existing publications do not always agree on the V gene identified, we report the same V gene for 8 out of 11 reports, and we report a member of the same V gene subfamily for the other 3 reports; and we report concordant J genes in 11 out of 11 reports. Furthermore, we report an identical HLA association for five of the previously published TCRs, with 1 sequence not significantly HLA-associated in this study.

**Table 2:**
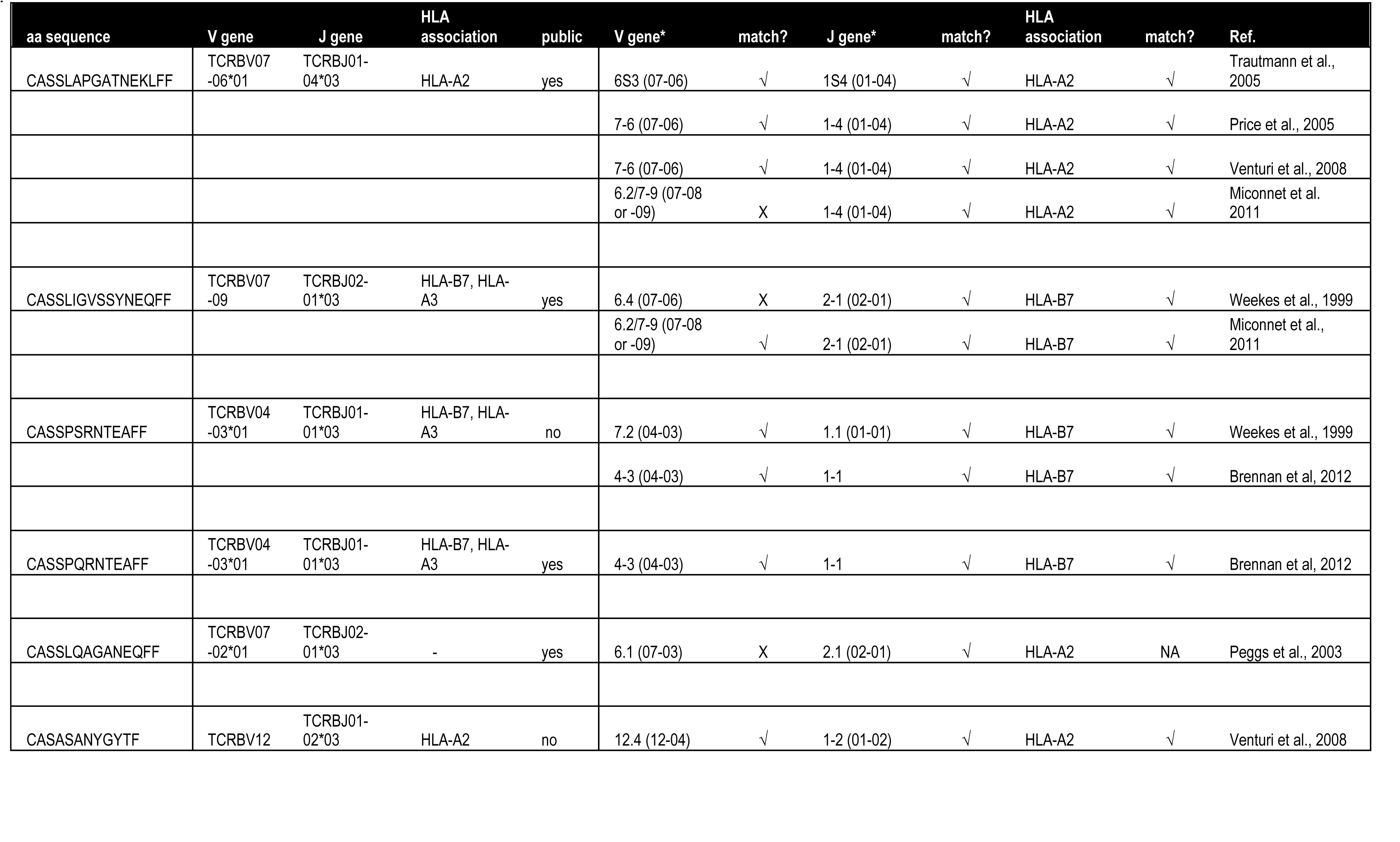
Concordance between this dataset and previously published data. The table lists the CDR3 amino acid sequence, V and J genes, and HLA association for each of the 6 CVM-associated TCRβ sequences identified in this study that had been previously reported as public, and compares these data to those from previous reports.

## DISCUSSION

We have demonstrated that information gleaned from rearranged T cell receptors can be used to infer disease status based on the presence of public T cell responses; the only requirement is a large sample of pathogen-positive and -negative samples with which to identify these public T cell responses. Because high-throughput sequencing of T cell receptors captures all T cell responses equally, and these store immunological memory to all pathogens in a common format, we believe that reading T cell memory by looking for known public responses will be a viable strategy for simultaneously diagnosing a wide range of infectious agents using a single peripheral blood sample and a simple, unified assay. More exploration will be needed to allow the application of the method to acute infections, given that T cell memory persists for years, and that we do not know how public clones will decay with time after an acute infection.

## ONLINE METHODS

### Experimental Cohort and Study Approval

Human peripheral blood samples were obtained from the Fred Hutchinson Cancer Research Center Research Cell Bank biorepository of healthy bone marrow donors. Donors underwent routine HLA-typing and CMV status typing at the time the samples were taken. Samples were obtained under a protocol approved and supervised by the Fred Hutchinson Cancer Research Center Institutional Review Board, following written informed consent.

### High-throughput TCRβ sequencing

Genomic DNA was extracted from peripheral blood samples using the Qiagen DNeasy Blood extraction Kit (Qiagen, Gaithersburg, MD, USA). We sequenced the CDR3 region of rearranged TCRβ genes, which was defined according to the IMGT collaboration ^28^. TCRβ CDR3 regions were amplified and sequenced using previously described protocols ^3^^,^^29^. Briefly, a multiplexed PCR method that uses a mixture of 60 forward primers specific to TCR Vβ gene segments and 13 reverse primers specific to TCR Jβ gene segments was employed. Reads of 87 bp were obtained using the Illumina HiSeq System. Raw HiSeq sequence data were preprocessed to remove errors in the primary sequence of each read, and to compress the data. A nearest neighbor algorithm was used to collapse the data into unique sequences by merging closely related sequences, to remove both PCR and sequencing errors.

### Identification of CMV-associated T cell receptors and classification of CMV status

On average, we identified 185,204 (+/– 84,171) unique TCRβ sequences for each of the 640 subjects, resulting in 83,727,796 unique TCRβ sequences in aggregate. Rather than attempting high dimensional CMV classification using all unique TCRβ sequences as potential features, a novel feature selection scheme was developed, which is described below in the “*Statistics”* section.

### Dimensionality reduction and machine learning

CMV memory burden was defined as the fraction of a subject’s unique TCRβ sequences that are CMV-associated (at a significance level defined by the procedure described above). This single dimension provided a strong discriminator between CMV+ and CMV- subjects (Figure 2C), enabling fast training of a one-dimensional logistic regression classifier of CMV status. Exhaustive leave-one-out cross validation (including recomputation of CMV- associated TCRβ sequences but conservatively assuming the same null distribution as in the slightly larger full dataset) was performed, and showed high accuracy across a broad range of p-value thresholds, with performance degrading at high FDR.

### Statistics

Since many TCRβ sequences are unique to a single subject (and consequently unique to either the CMV+ or CMV– classes), it was vital to control false discovery rate in feature selection to avoid overfitting to the many spurious associations of unique TCRβ sequences with CMV status. Each unique TCRβ rearrangement, identified by both the V and J gene assignment and the CDR3 amino acid sequence, was tested for CMV association by subjecting it to a one-tailed Fisher exact test for its incidence in CMV– and CMV+ subjects. Specifically, letting *n*_*ij*_ denote the number of subjects with CMV status *j* (with *j* – or +) and TCRβ sequence *i* present, we compute a p-value *p*_*i*_ by performing Fisher’s exact test on the following contingency table where *N*_+_ and *N*_–_ denote the total number of subjects positive or negative for CMV:

|  | **CMV+** | **CMV-** |
| --- | --- | --- |
| **TCRβ sequence $i$ present** | $n_{i+}$ | $n_{i-}$ |
| **TCRβ sequence $i$ not present** | $N_+ - n_{i+}$ | $N_- - n_{i-}$ |

To characterize a rejection region in the presence of many weakly dependent hypotheses (one for each unique TCRβ sequence), we randomly permuted CMV status assignments 100 times. In each permutation, we recomputed a p-value for each TCRβsequence and recorded the number of rejections at the nominal p-value threshold. Approximating the total fraction of true null hypotheses as 1 in these permutations, this allowed us to estimate the false discovery rate (FDR) as the ratio of the mean number of rejections under permutation to the actual number of rejections ^30^.

## ACKNOWLEDGEMENTS

J.G. and J.A.H. obtained the DNA samples and determined CMV status and HLA type of the subjects, R.O.E., C.S.C., M.R. and H.S.R. conceived and designed the experiments, M.R. generated the sequence data, R.E.O, W.S.D., M.V. and C.D. analyzed the results, R.O.E. and W.S.D. performed the statistical analyses, M.V. and C.D. performed the literature searches of CMV-associated TCR clones, and R.O.E., W.S.D, M.V. and H.S.R. wrote the manuscript.

## Conflict of Interest

H.S.R. and C.S.C. have consultancy, equity ownership, patents and royalties with Adaptive Biotechnologies; R.O.E., W.S.D., M.V., C.D and M.R have employment and equity ownership with Adaptive Biotechnologies.

